# Smell what you hardly see: Odors assist categorization in the human visual cortex

**DOI:** 10.1101/2021.05.25.445626

**Authors:** Diane Rekow, Jean-Yves Baudouin, Karine Durand, Arnaud Leleu

## Abstract

Visual categorization is the brain ability to rapidly and automatically respond to widely variable visual inputs in a category-selective manner (i.e., distinct responses between categories and similar responses within categories). Whether category-selective neural responses are purely visual or can be influenced by other sensory modalities remains unclear. Here, we test whether odors modulate visual categorization, expecting that odors facilitate the neural categorization of congruent visual objects, especially when the visual category is ambiguous. Scalp electroencephalogram (EEG) was recorded while natural images depicting various objects were displayed in rapid 12-Hz streams (i.e., 12 images / second) and variable exemplars of a target category (either human faces, cars, or facelike objects in dedicated sequences) were interleaved every 9^th^ stimulus to tag category-selective responses at 12/9 = 1.33 Hz in the EEG frequency spectrum. During visual stimulation, participants (N = 26) were implicitly exposed to odor contexts (either body, gasoline or baseline odors) and performed an orthogonal cross-detection task. We identify clear category-selective responses to every category over the occipito-temporal cortex, with the largest response for human faces and the lowest for facelike objects. Critically, body odor boosts the response to the ambiguous facelike objects (i.e., either perceived as nonface objects or faces) over the right hemisphere, especially for participants reporting their presence post-stimulation. By contrast, odors do not significantly modulate other category-selective responses, nor the general visual response recorded at 12 Hz, revealing a specific influence on the categorization of congruent ambiguous stimuli. Overall, these findings support the view that the brain actively uses cues from the different senses to readily categorize visual inputs, and that olfaction, which is generally considered as poorly functional in humans, is well placed to disambiguate visual information.

## Introduction

Vision is commonly considered the dominant sense in humans whereas olfaction is deemed as poorly functional. This is illustrated by the fact that nearly 3 out of 4 persons are more afraid of blindness than of any other sensory deprivation, while no one report anosmia as the scariest deprivation (Hutmacher, 2019). The function of olfaction has long been confined to alertness (Herrick, 1933), as most of our chemical environment remains unnoticed (Sela and Sobel, 2010), and we are consistently better at *detecting* than *identifying* an odor (Cain, 1979; Yeshurun and Sobel, 2010). In fact, odor recognition appears rather undetermined and flexible (Barwich, 2019; Cain, 1979), and is largely influenced by contextual cues such as colors (e.g., Morrot et al., 2001; Zellner et al., 1991) or verbal labels (Herz and von Clef, 2001). As a result, olfactory-visual interactions have often been investigated through the lens of vision modulating odor perception (e.g., Gottfried and Dolan, 2003; Jadauji et al., 2012; Manesse et al., 2020).

Over the last decades, it has yet been progressively established that humans possess a keen sense of smell (McGann, 2017; Schaal and Porter, 1991), and mounting evidence reveals how olfaction influences other sensory modalities, in particular vision. Odors indeed attenuate the attentional blink for congruent visual objects (Robinson et al., 2013), help color recognition if odors and colors were previously paired (Demattè et al., 2006), improve congruent object detection in visual scenes (Seigneuric et al., 2010; Seo et al., 2010), and bias perception towards the congruent object during binocular rivalry (Zhou et al., 2010). At the neural level, odor (in)congruency modulates the scalp electroencephalographic (EEG) activity elicited by a visual stimulus (Ohla et al., 2018), or by an auditory cue that follows paired odor and visual stimuli and signals that the visual stimulus must be explicitly categorized (Hörberg et al., 2020). Olfactory-visual integration activates a broad neural network (Ripp et al., 2018), including the lingual and fusiform gyri (traditionally considered as visual brain regions), which respond as a function of the reported congruency between a visual object and an odor (Lundström et al., 2019).

The influence of odors on vision has also been extensively described for one of the most important objects of the human visual environment, i.e., faces. Odors facilitate face memory (Cecchetto et al., 2020; Steinberg et al., 2012) and orient judgments of face attractiveness (Demattè et al., 2007; Parma et al., 2012; Rikowski and Grammer, 1999), face sex (Kovács et al., 2004), or face-evoked personality traits (Cook et al., 2015, 2017, 2018; Dalton et al., 2013; Li et al., 2007). Emotional body odors (i.e., collected in anxiogenic or happy contexts) elicit (in)congruency effects on the perception of facial expressions (Kamiloğlu et al., 2018; Mujica-Parodi et al., 2009; Rocha et al., 2018; Wudarczyk et al., 2016; Zernecke et al., 2011; Zhou and Chen, 2009), which is also biased by hedonically-contrasted non-body odors (Cook et al., 2017; Leleu et al., 2015a; Leppänen and Hietanen, 2003; Seubert et al., 2010; Syrjänen et al., 2017, 2018). The neural underpinnings of these odor influences on facial information have been explored, revealing various patterns of modulations in “visual” brain regions (Cecchetto et al., 2020; Wudarczyk et al., 2016; Novak et al., 2015; Seubert et al., 2010), or in the EEG activity elicited by the face stimulus (Adolph et al., 2013; Cook et al., 2017; Forscher and Li, 2012; Leleu et al., 2015b; Poncet et al., 2021; Rubin et al., 2012; Syrjänen et al., 2018). Interestingly, given the high relevance of the sense of smell at the beginning of life (Schaal et al., 2020, for review) compared to the relative immaturity of the visual system (Braddick and Atkinson, 2011), odors strongly influence how infants look at faces (Durand et al., 2020, 2013), or how their brain responds to facial information (Jessen, 2020; Leleu et al., 2020; Rekow et al., 2020, 2021b).

Despite consensual evidence that odors influence visual perception, several important questions remain unanswered. *First*, whether odors are truly able to influence neural visual categorization is unclear. Visual categorization is the brain ability to rapidly (i.e., at a glance) and automatically (i.e., without volitional control) respond to a certain class of visual objects (e.g., Bugatus et al., 2017; Thorpe et al., 1996), relying on a set of category-selective regions in the ventral occipito-temporal cortex (VOTC). These regions go beyond visual inputs and generate categorical responses, that is, distinct responses to different categories (i.e., between-category discrimination) and similar responses to different exemplars of one category despite their physical variability (i.e., within-category generalization, Bracci and Op de Beeck, 2016; Hagen et al., 2020; Jacques et al., 2016b). While well-known category-selective visual regions (e.g., the fusiform gyrus) have been associated with odor effects in previously reviewed neuroimaging studies (Cecchetto et al., 2020; Lundström et al., 2019; Seubert et al., 2010; Wudarczyk et al., 2016), their activity was often considered for different behavioral responses to a single category rather than different visual categories irrespective of the behavioral output (i.e., to measure a category-selective neural response). Similarly, in EEG studies, odor effects have been rarely explored for selective responses to a variety of inputs from a given category contrasted to many other object categories (e.g., only a few individual faces for each emotion in the numerous studies investigating the effect of odors on the perception of facial expressions) and sometimes measured at late latencies over parietal and frontal regions (e.g., Hörberg et al., 2020; Ohla et al., 2018), contrary to occipito-temporal category-selective EEG responses (e.g., Jacques et al., 2016a). To our knowledge, the only EEG studies measuring how odors modulate a category-selective visual response have been conducted in 4-month-old infants (Leleu et al., 2020; Rekow et al., 2020, 2021b). Whether odors affect automatic visual categorization in the adult brain is still to be established.

*Second*, since both faces and body odors convey a wealth of information about our conspecifics (e.g., identity, sex, age) and their internal states (e.g., emotion, health), prior interest for odor-face integration focused on person-related information. However, for faces, the initial categorization level is the mere categorization of a visual object as a face before categorizing fine-grained information such as identity or facial expression (Quek et al., 2020). As far as we know, no study has addressed whether a body odor may tune this generic face categorization function in adults. At 4 months of age, maternal body odor orients the infant’s gaze towards a face when it is paired with a car (Durand et al., 2013), and enhances a face-selective neural response over the right occipito-temporal cortex (Leleu et al., 2020). The latter effect is selective to face categorization, as no effect is found for a nonface category (i.e., cars: Rekow et al., 2020), except for nonface objects perceived as faces (i.e., facelike objects; Rekow et al., 2021b). Hence, whether olfactory-visual interaction for generic face categorization is maintained in adulthood must be examined.

*Third* and finally, considering that the adult visual system readily categorizes faces and other visual objects from the sole visual input in typical conditions, whether the putative odor influence on category-selective responses in the adult brain depends on the ambiguity of the visual input has to be delineated. Indeed, odor effects on category-selective neural responses in 4-month-old infants (Leleu et al., 2020; Rekow et al., 2020, 2021b) have been observed when the visual system is still immature and visual experience is poor compared to that of adults. In addition, the strength of the odor effect for a given infant increases as the sole visual input is weakly effective in evoking a response (Rekow et al., 2021b), in line with the inverse effectiveness principle of multisensory integration (Regenbogen et al., 2016; Stein and Meredith, 1993). Similarly, numerous adult studies found the largest odor effects on facial expression recognition for ambiguous stimuli (Forscher and Li, 2012; Leleu et al., 2015a; Mujica-Parodi et al., 2009; Novak et al., 2015; Rubin et al., 2012; Zernecke et al., 2011; Zhou and Chen, 2009). Accordingly, whether congruent odors act as disambiguating cues on category-selective visual responses in the adult brain (i.e., the so-called disambiguation function of multisensory integration; Ernst and Bülthoff, 2004) has to be explored.

Here we address these outstanding issues using an EEG frequency-tagging approach. We focus on the visual categorization of human faces, nonface objects resembling faces (i.e., facelike objects), and cars contrasted to a variety of other living and non-living objects. Natural images were displayed at a fast rate of 12 Hz (i.e., 12 images / s) and the target category (i.e., human faces, facelike objects or cars in different sequences) was inserted every 9th stimulus (i.e., at 1.33 Hz) in 24-second-long sequences while participants performed an orthogonal cross-detection task. We isolate a general visual response common to all stimuli at 12 Hz and harmonics (i.e., integer multiples) in the EEG frequency spectrum, and, most importantly, a category-selective response at 1.33 Hz and harmonics (Jacques et al., 2016a; Rossion et al., 2015; Norcia et al., 2015 for review). The latter response is a direct differential response to the target category (i.e., reflecting its discrimination from the other categories displayed in the sequence that generalizes across the various exemplars of the target category) generated by category-selective regions in the VOTC (Gao et al., 2018; Hagen et al., 2020; Jonas et al., 2016). Thanks to the fast rate of stimulation and the orthogonal behavioral task, the category-selective response measures single-glance and automatic visual categorization. During visual stimulation, participants were alternatively and blindly exposed to a body, a gasoline or a baseline (i.e., mineral oil) odor context, the two formers being chosen for their expected congruency with face(like) and car stimuli, respectively. Following previous studies in infants (Leleu et al., 2020; Rekow et al., 2020, 2021b), and since faces and cars are readily categorized in the adult brain whereas facelike objects are more ambiguous (i.e., they elicit a lower response than genuine human faces and are reported by only a fraction of participants in such a rapid and implicit mode of visual stimulation; Rekow et al., 2021a), we hypothesized that the congruent body odor context mainly enhances the visual categorization of facelike objects, this effect depending on the perceptual awareness of facelike objects (i.e., face pareidolia).

## Materials and methods

### Data availability

Access to the data will be granted to named individuals. Specifically, requestors must complete a formal data sharing agreement to obtain the data from the corresponding author.

### Participants

Twenty-six participants (14 females, 4 left-handed, mean age ± SD: 25 ± 4.5 years old) were recruited and compensated for their participation. All were healthy at the time of the study and reported normal or corrected-to-normal vision, and no history of allergy, sensory impairment, psychiatric or neurological disorder. The work described has been carried out in accordance with The Code of Ethics of the World Medical Association (Declaration of Helsinki) for experiments involving humans. Participants provided written informed consent prior to beginning. A full debriefing after the experiment explained the whys and whereabouts of the study and revealed they have been exposed to implicit olfactory stimulation during testing. An additional consent was thus obtained after full disclosure.

### Visual stimuli

We used 368 natural images of objects unsegmented from their background (examples in Figure 1A) divided in 4 subsets: 66 human faces (33 females), 66 cars, 66 facelike stimuli (i.e., nonface objects inducing face pareidolia) and 170 base objects of numerous living and non-living categories (e.g., plants, vegetables, animals, man-made objects), some of them being common with facelike stimuli. These stimuli have been used in previous experiments using an analogous approach in infants (Leleu et al., 2020; Rekow et al., 2021b, 2020) and adults (e.g., Quek et al., 2018; Rekow et al., 2021a). Each image contained a single item, depicted off-centered in the image to increase physical variability across category exemplars. In addition, items varied in size, viewpoint, lighting condition and background. After being cropped to a square, they were resized to 300 × 300 pixels. Displayed on a screen at a 57-cm distance, they roughly covered 8.3° of visual angle.

**Figure 1.**
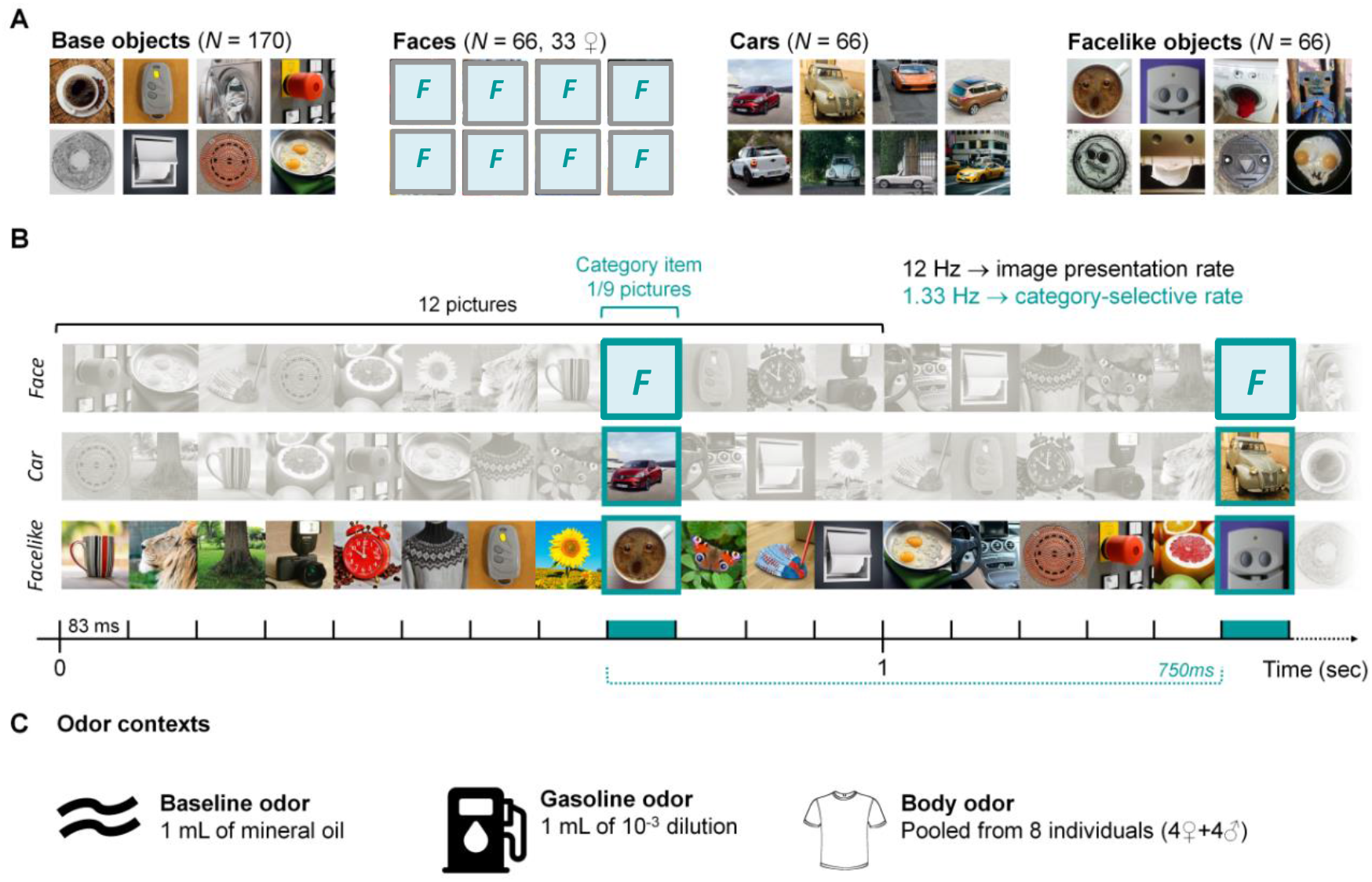
Stimuli and experimental paradigm. **A**. Examples of variable unsegmented images used as stimuli and depicting base objects (*N* = 170), human faces (*N* = 66, 33 females, replaced by F placeholders for the preprint version), cars (*N* = 66) and facelike objects (*N* = 66). **B**. Excerpt of ≈ 1.5s (out of 24) of visual stimulation at a 12-Hz rate of image presentation (i.e., 12 images per second, 83 ms per image). Base objects are presented while faces, cars or facelike objects (in different sequences) are inserted every 9^th^ stimulus (i.e., at a 1.33-Hz category-selective rate, 750 ms between two exemplars). **C**. Odor contexts (baseline, gasoline and body odor) are presented throughout each sequence (one odor per sequence).

### Odor stimuli

Three odor contexts were used: a generic human body odor (i.e., pooled across 8 donors), a gasoline odor, and a baseline odor (i.e., scentless mineral oil). The two formers were chosen for their congruency with the visual categories (i.e., face(like)s and cars) and the latter as a control odor condition. Pilot experiments were conducted to characterize the sensory properties of body odors and match the gasoline odor with them (see Supplementary Methods for details).

The body odor consisted in axillary sweat samples collected on cotton pads. They were collected from 16 independent non-smokers donors (8 females, mean age ± SD: 25 ± 4 years old). All donors followed a 24-hour hygiene procedure (see Supplementary Methods for details). Two pools of 8 individuals each (4 females) were created by matching sampling duration and age across pools (see Table S1). These pools were used for a pilot odor evaluation (see Tables S2, S3 and Supplementary Methods) and the main EEG experiment (see below). The gasoline odor consisted in 1 mL of 10^−3^ gasoline oil diluted in mineral oil and disposed on a volume of cotton pads equivalent as for the body odor condition (i.e., 1 cotton pad, cut in 16 units). The odorless baseline odor consisted in a cotton pad impregnated with 1 mL of scentless mineral oil and cut in 16 units. Odors for each participant were prepared before testing in a separate room: units of cotton pads containing body odor, gasoline or mineral oil were put in dedicated 60 mL sealed glass flasks and left at room temperature (+20°C).

### Procedure

EEG was recorded in a sound and light-attenuated cabin equipped with an air-vacuum. To reduce additional olfactory noise, the non-smoker experimenter used scentless soap and avoided consuming any odorant product prior to testing. In the cabin, participants were seated at a 57-cm distance from the screen with their head on a chinrest. The screen (24-inch LED) displayed images with a refresh rate of 60 Hz and a resolution of 1920 × 1080 pixels on a uniform grey background (i.e., 128/255 in greyscale). To diffuse the odorants, we used an odor-delivering device adapted from previous studies (Leleu et al., 2015b; Poncet et al., 2021). The three odor flasks were connected to a device delivering a constant flow of scentless air originating from a tank of pressured air purified by charcoal filters and set at room temperature. The airflow was delivered at an undetectable pressure (i.e., 0.5 bar) to avoid the mechanical sensation of air on the skin and to ensure unawareness of olfactory stimulation throughout the experiment. The airflow was directed to one of the three flasks by a hand-activated valve from where a tube was connected to the chinrest to diffuse odors directly under the nose of the participants in the cabin. The flasks and the odor diffusing system were hidden from the participants.

We used an EEG frequency-tagging approach (Norcia et al., 2015) to measure rapid (i.e., single-glance) and automatic (i.e., without explicit intention) visual categorization in the brain. The design was adapted from previous studies which successfully isolated category-selective responses at different levels of brain organization in adults (e.g., Gao et al., 2018; Hagen et al., 2020; Jacques et al., 2016a), and infants (de Heering and Rossion, 2015; Leleu et al., 2020; Rekow et al., 2021b, 2020). Base objects were presented without inter-stimulus interval at a rapid 12-Hz rate (i.e., 12 images / second, ≈ 83 ms per image, Figure 1B) and images of either human faces, cars, or facelike objects (one target category per sequence) were periodically interspersed every 9^th^ stimulus, corresponding to a category-selective rate of 1.33 Hz (i.e., 12 / 9; 750 ms between each category exemplar). With this approach, we isolate two distinct brain responses in the EEG frequency spectrum: (1)a general visual response at 12 Hz and harmonics (i.e., integer multiples) elicited by the information rapidly changing 12 times per second (e.g., local contrast) and (2)a category-selective response at 1.33 Hz and harmonics reflecting the visual categorization of the target category. The latter response is elicited by populations of neurons that selectively respond to this category in the VOTC (Gao et al., 2018; Hagen et al., 2020; Jonas et al., 2016).

The visual stimulation sequences were as follows: After a fixed interval of 1.5 seconds, a fade-in ramping from 0 to 100% contrast depth lasted 1.417 s before 23.333 s of full-contrast stimulation. A 0.667-s fade-out of decreasing contrast followed and the sequence closed on a 0.083 s of post-stimulation interval of grey background. For the target categories, each set of 66 images was randomly divided into two 33 stimuli sets, each set being used in a single sequence. All base objects were used in every sequence. During each sequence, stimuli were randomly selected from their respective sets. For the body odor, each participant was exposed to only one body odor pool (see Supplementary Methods). Given the high volatility of the gasoline odor, two 1 mL samples were presented each for one half of the experiment. The nine experimental conditions were presented 4 times each: 3 odor contexts (body, gasoline, baseline) × 3 visual categories (human faces, facelike objects, cars) × 4 repetitions (2 subsets of stimuli presented twice). Each participant was thus tested for 36 sequences organized in 12 blocks of 3 sequences. In each block, odor conditions were paired each with one visual category, such that every odor and visual conditions were presented once within a block. These odor-visual associations were alternated between blocks.

After ensuring the participant was still and ready, the experimenter started odor diffusion and launched visual sequences. Odor diffusion started 5 seconds before each visual sequence and remains for the whole sequence. A minimum interval of 25 seconds was introduced between visual sequences and during which the baseline odor was diffused. In other words, at the end of each visual sequence, the experimenter immediately replaced the odor stimulation by the baseline odor if necessary and waited 25 seconds before asking the participant if they were ready for the next sequence.

To ensure that participants stayed focus on the visual stimulation, they performed an orthogonal behavioral task consisting in the detection of a 250 × 250 pixel-large white cross (3-pixel thick, 200 ms duration) superimposed on the images at the center of the screen. The cross appeared randomly six times per sequence, with a 2-second-minimum interval between appearances. Participants were instructed to press the spacebar (simultaneously with both index fingers) as rapidly as possible when they detect the cross. An ANOVA was run on accuracy and response times for correct detections, and revealed no effect of *Category* (face, car, facelike), *Odor* (body, gasoline, baseline), and *Category* × *Odor* interaction (all *F*s < 1). The mean accuracy was near ceiling (97.7 ± 0.3 (*SD*) %) with a mean response time of 396 ± 28 (*SD*) ms.

After the EEG experiment, participants were asked to fill a questionnaire intended to document (1) the non-detection of odors during the experiment and their evaluation, (2) naivety regarding the frequency-tagging approach and the tagged categories, and (3) whether they perceived the facelike stimuli (see Rekow et al., 2021a, for details). No participants declared having noticed the presence of the airflow and the diffusion of odors during the experiment, nor the periodicity of the presentation, or the dissociation of sequences based on target categories. A total of 9 participants (i.e., 35%) declared having perceived the facelike stimuli on the course of the experiment; they will be henceforth designated as perceptually “aware” participants vs. “unaware” participants for those who did not notice the facelike objects (i.e., the 17 remaining participants). After the experimenter disclosed the diffusion of odors, participants were asked to rate the odorants (see Supplementary Methods, Tables S2 and S3). Gasoline and body odors did not differ in perceived pleasantness, intensity and familiarity (all *t*s < 1.95, all *p*s > .06).

### EEG recording and preprocessing

Scalp electroencephalogram (EEG) recording started once the participant was installed in the cabin. It was continuously acquired until the end of the experiment. A 64-channel Biosemi Active-Two amplifier system was used, with Ag/AgCl electrodes disposed according to the 10–10 classification montage (BioSemi, The Netherlands) and sampled at 1024 Hz. Reference and ground were constituted by the active electrode CMS (Common Mode Sense) and the passive electrode DRL (Driven Right Leg), respectively. Electrode offset was set below ± 15 μV for all electrodes.

Following EEG analyses were run on Letswave 6 (https://www.letswave.org/) implemented on Matlab 2017 (MathWorks, USA). Continuous individual datasets were first highpass filtered at 0.1 Hz using a 4^th^-order Butterworth filter, then resampled to 200 Hz. Epochs were segmented from the start of the fade-in until 0.583 ms after the end of fade-out (i.e., for 26 s) resulting in 36 segments per participant. To identify eye-blinks and additional high (i.e., > 200µV) artifacts over frontal or temporal electrodes, an Independent Component Analysis (ICA) using a square mixing matrix was computed for each epoch and participant. The mean ± SD number of ICs removed was 4 ± 2 (range: 1–8). Additional artifact-ridden electrodes were linearly interpolated from 3 to 5 (depending on edge/central locations) immediately neighboring channels, for an average of 2 ± 2 interpolations per participant (range: 0–7). Epochs were then re-referenced to the average of the 64 channels.

### EEG frequency-domain analysis

EEG data analysis was largely similar to previous frequency-tagging studies on visual categorization (Jacques et al., 2016a; Rekow et al., 2021a; see Retter and Rossion, 2016 for a discussion). Epochs were precisely segmented to comprise an exact number of category-selective 1.33-Hz cycles, i.e., into 24-s-long epochs, starting from the end of the fade-in (i.e., first target category exemplar) to the end of the fade-out, for a total of 32 cycles. To reduce neural activity non-phase-locked to the presentation of the target stimuli, epochs were then individually averaged for the 4 repetitions of each condition, resulting in 9 epochs of 24 s per participant (i.e., 1 per experimental condition). A fast Fourier transform (FFT) was applied to every epoch and amplitude spectra were extracted for all channels with a high frequency resolution of 1/24 = 0.0417 Hz.

Next, we evaluated the number of harmonics to retain for having a thorough estimation of each brain response. To consider an identical number of harmonics across experimental conditions, individual data were grand-averaged across odor contexts and visual categories, and channels were pooled together. *Z*-scores were computed on amplitude spectra as the difference between each frequency bin and the mean surrounding noise estimated from the 20 adjacent bins (10 on each side) excluding the most extreme (minimum and maximum) and immediately adjacent bins, divided by the standard deviation of the noise. Harmonics were considered until *Z*-scores ceased to be consecutively significant (*Z* > 1.64, *p* < .05, one-tailed, signal > noise). For the category-selective response, harmonics were significant until the 14^th^ harmonic (i.e., 18.67 Hz). For the general visual response, harmonics were significant until the 4^th^ harmonic (i.e., 48 Hz; harmonics above the 50-Hz response elicited by AC power were not considered). To provide a summary representation of both responses (Retter and Rossion, 2016), they were compiled by summing significant harmonics (excluding the 12-Hz harmonic (i.e., general response) for the category-selective response) for each condition, channel and participant. In the following sections, mentions of both responses will refer to these overall responses summed across harmonics.

The magnitude of each brain response was quantified by a baseline-corrected amplitude value expressed in microvolt (µV) obtained by subtracting the mean background noise from the raw amplitudes, based on the same noise estimation as defined above. Considering that each visual category may recruit different neural populations, we defined regions of interest (ROIs, Figure S1) separately for each category from group-level data. Baseline-corrected amplitudes at each electrode were ranked from highest to lowest (Tables S4 and S5). For the three category-selective responses, the six best electrodes were P10, PO8, P8, P9, PO7 and P7 (different order for each visual category; Table S4). Two ROIs were thus considered over the right occipito-temporal cortex (rOT) and the left occipito-temporal cortex (lOT) to account for putative hemispheric asymmetries. For all three visual categories, a single ROI was built for the general visual response over the middle occipital cortex (4 best channels: O1/2, Oz, Iz). For both brain responses, ROIs were used for statistical analyses.

Statistical analyses were computed separately for each brain response. The significance of each brain response at both group and individual levels was estimated using *Z*-scores (see above) calculated on uncorrected amplitudes. Repeated-measures ANOVA were also run on individual baseline-corrected amplitudes. For the category-selective response, *Odor* (body, gasoline, baseline), *Category* (faces, cars, facelike objects) and *Hemisphere* (rOT, lOT) were used as within-subject factors, and *Group* (aware, unaware) as a between-subject factor. For the general visual response, the same factors were considered without the factor *Hemisphere* (only one ROI). Mauchly’s test for sphericity violation was computed and Greenhouse-Geisser correction (*ε*) for degrees of freedom was applied whenever sphericity was violated. Effect sizes are reported with partial eta squared (*η*_*p*_^*2*^). For significant *Odor* effects, orthogonal contrasts were calculated to qualify the effects. Since the amplitude of the category-selective response can be highly different between visual categories (Jacques et al., 2016a; see Rekow et al., 2021a for the difference between the face- and facelike-selective responses), the *Odor* effect on the weakest category-selective response might be masked by the largest responses in the omnibus ANOVA. Hence, we also ran a repeated-measures ANOVA after having normalized each category-selective response by its overall amplitude over the scalp (McCarthy and Wood, 1985). A significant *Odor* effect for a given visual category was then further explored by directly conducting a repeated-measures ANOVA on the difference between odor conditions (expressed in non-normalized baseline-corrected amplitudes) for this specific category, and *Z*-scores (see above) were calculated to estimate the significance of the effect (*Z* > |1.96|, *p* < .05, two-tailed, effect ≠ 0).

## Results

### Neural categorization of human faces, cars, and facelike objects across odor contexts

Despite the high constraints put on the visual system to categorize human faces, cars, and facelike objects at a glance within 12-Hz streams of numerous living and non-living objects, the three visual categories elicit a clear selective response (i.e., a direct differential response that generalizes across category exemplars) distributed on several harmonics (i.e., 1.33 Hz and integer multiples) in the EEG frequency spectrum, especially over the occipito-temporal cortex (Figure 2A). Summed across harmonics and averaged across hemispheres, every response is significant (*Z* = 21.1, 11.3, and 2.13 respectively for the face-, car-, and facelike-selective responses, all *p*s < .017). After noise correction, the face-selective response appears as the largest (mean amplitude across hemispheres ± *SEM*: 2.56 ± 0.21 µV), followed by the car-selective response (1.38 ± 0.12 µV, 54% of the face-selective response) and the facelike-selective response (0.29 ± 0.05 µV, 21% of the car-selective response, 12% of the face-selective response), as revealed by a main effect of *Category* (*F* (1.6, 39.8) = 71.1, *ε* = 0.80, *p* < .001, *η*_*p*_^*2*^ = 0.74). Accordingly, while every participant presents with a significant (i.e., *Z* > 1.64) face-selective response and 25 participants out of 26 with a significant car-selective response, the facelike-selective response is significant in only 14 participants out of 26. In addition, the three responses are larger over the right hemisphere (rOT > lOT, main effect of *Hemisphere*: *F* (1, 25) = 9.39, *p* = .005, *η*_*p*_^*2*^ = 0.27; Figure 2B), especially the facelike-selective response (0.36 ± 0.08 µV vs. 0.23 ± 0.05 µV, i.e. +59% over rOT; faces: 2.88 ± 0.29 µV vs. 2.25 ± 0.22 µV, +28% over rOT; cars: 1.61 ± 0.19 µV vs. 1.16 ± 0.09 µV, +39% over rOT).

**Figure 2.**
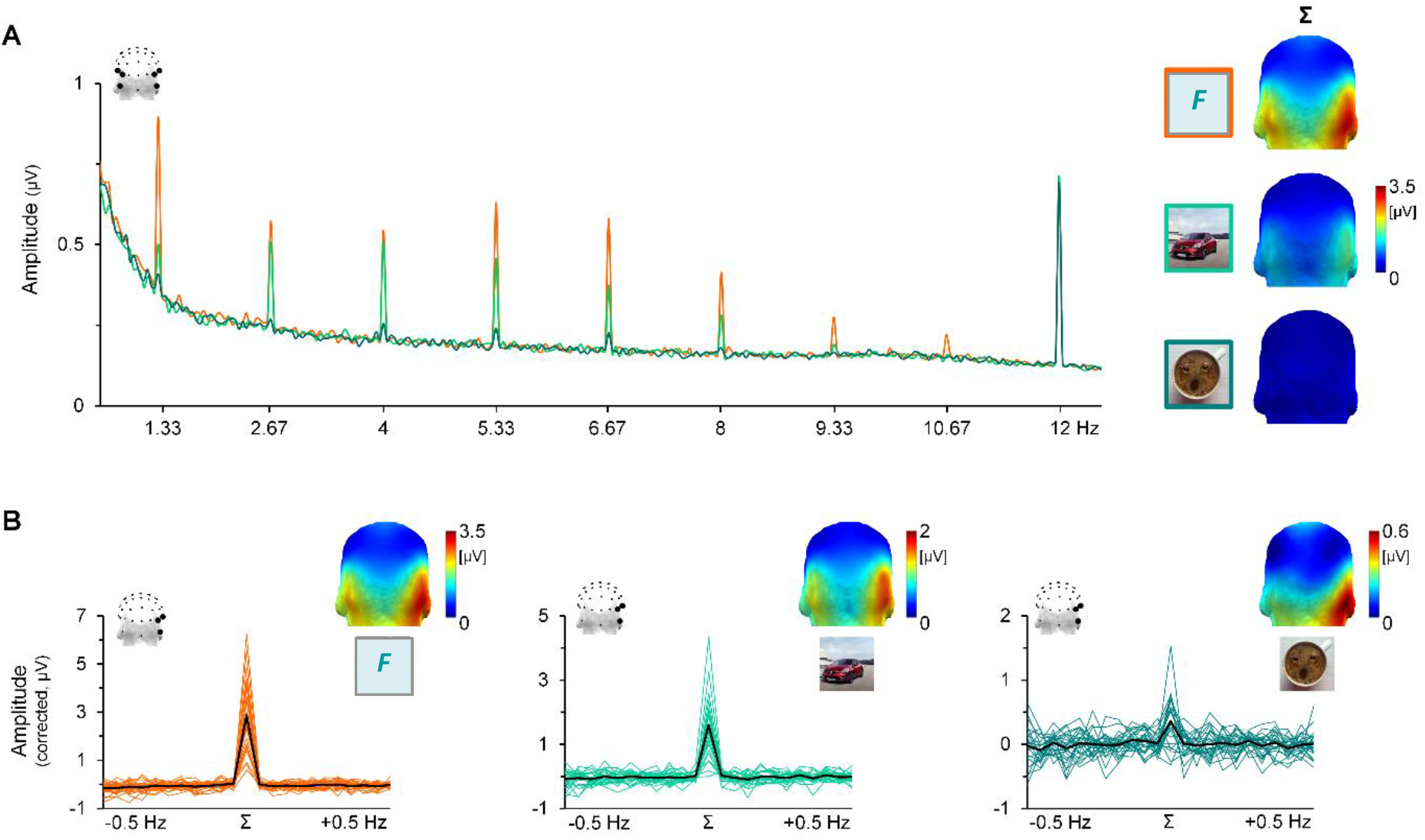
EEG frequency spectrum averaged across odor contexts for each visual category. **A. Left:** Grand-averaged FFT amplitude spectra (uncorrected) recorded for sequences presenting human faces (orange, replaced by F placeholders for the preprint version), cars (light green) and facelike objects (dark green) among base objects. All types of sequences elicit clear responses (larger than surrounding frequencies) at the 12-Hz frequency of stimulation and at the 1.33-Hz category-selective frequency and its harmonics (i.e., integer multiples, here displayed from 2.67 Hz to 10.67 Hz) over bilateral occipito-temporal channels (P9/10, PO7/8 and P7/8). **Right:** 3D head-maps (back view, same scale) showing the topography and the magnitude (in baseline-corrected amplitude) of each category-selective response summed across harmonics (Σ). **B**. Baseline-corrected amplitude of the category-selective responses summed across significant harmonics (Σ) compared to surrounding frequencies (± 0.5 Hz, baseline-corrected amplitude ≈ 0, signal ≈ noise) over the right occipito-temporal region (rOT). The black line depicts the mean of the group and colored lines represent individual responses. Adjusted-scale 3D head-maps (back view) are shown for each category.

### Category-selective responses as a function of odor context

Visual inspection suggests that, compared to the other odor contexts, the body odor context increases the facelike-selective response, whereas both the face- and car-selective responses seem identical across odors (Figure 3). However, given the very low amplitude of the facelike-selective response compared to the two other responses, the omnibus ANOVA did not reveal any significant interaction involving the *Category* and *Odor* factors (all *F*s < 1.35, all *ps* > .26). We therefore conducted another ANOVA with the same factors after having normalized the responses by their whole-scalp amplitude (McCarthy and Wood, 1985) to equate their magnitude, and found a significant *Category* × *Odor* interaction (*F* (2.3, 55.8) = 4.47, *ε* = 0.58, *p* = .012, *η*_*p*_^*2*^ = 0.16). As suggested by visual inspection, the *Odor* effect is significant for the facelike-selective response (*F* (2, 48) = 5.12, *p* = .009, *η*_*p*_^*2*^ = 0.18), while non-significant for both the face-selective and car-selective responses (all *F*s < 1). For the facelike-selective response, a significant difference between the body odor context and the two other contexts (*F* (1,24) =9.58, *p* = .005, *η*_*p*_^*2*^ = 0.29) explains 70% of the effect. The remaining difference between the baseline and gasoline odors is not significant (*F* (1,24) = 2.48, *p* = .13, *η*_*p*_^*2*^ *=* 0.09).

**Figure 3.**
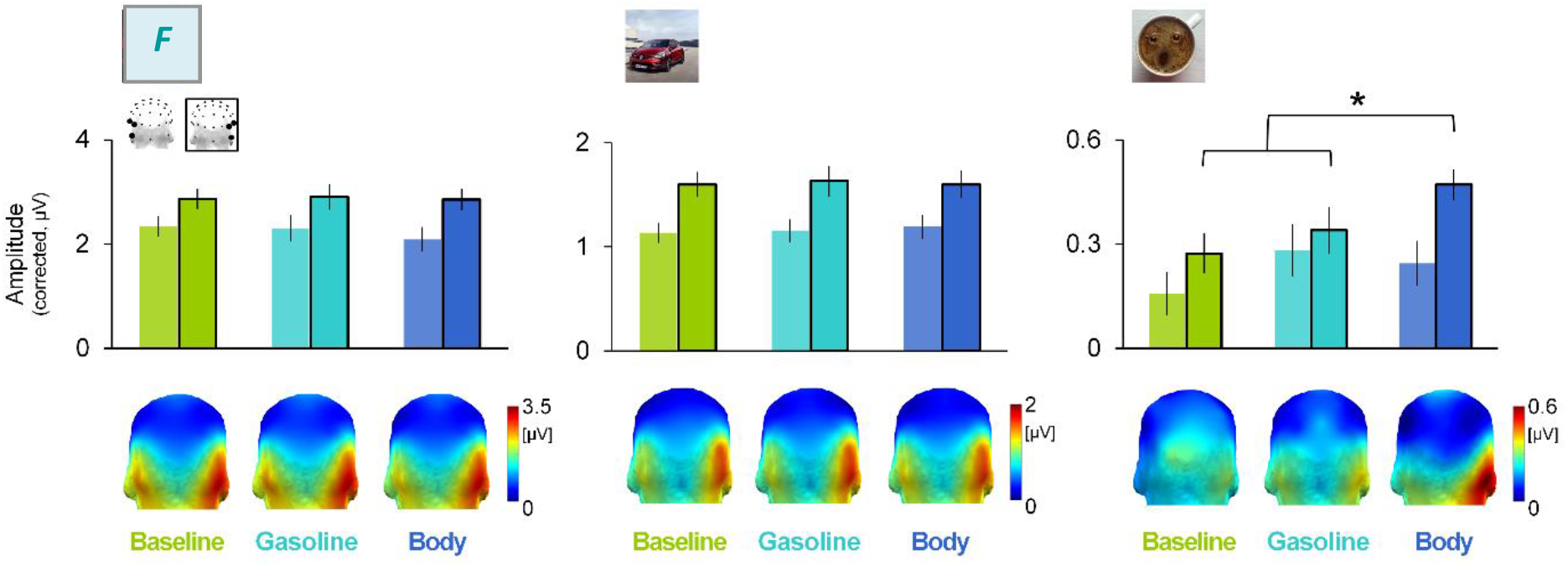
Category-selective responses according to odor context. Summed baseline-corrected amplitudes of the category-selective responses for each visual category (left: human faces, replaced by F placeholders for the preprint version, middle: cars, right: facelike objects), odor context (green: baseline odor, light blue: gasoline odor, dark blue: body odor) and hemisphere (unframed: lOT, framed: rOT) together with corresponding 3D head-maps (back view, adjusted scales). Error bars represent standard errors of the mean; *: *p* < .05.

Descriptively, the facelike-selective response (Figure 3, right panel) is particularly increased by the body odor context over the right hemisphere (rOT), with a ≈ 54% larger amplitude in this odor context than in the two other contexts (0.47 ± 0.09 µV vs. 0.27 ± 0.09 and 0.34 ± 0.11 µV for the baseline and gasoline odors, respectively). In contrast, the response is less variable over lOT, its amplitude ranging from 0.16 ± 0.06 to 0.28 ± 0.07 µV, the latter being observed for the gasoline odor context. As a result, we dissociated the *Odor* effect as a function of the hemisphere and found a significant *Odor* effect over rOT (*F* (2, 48) = 3.71, *p* = .032, *η*_*p*_^*2*^ = 0.13), but not lOT (*F* (2, 48) = 1.32, *p* = .28, *η*_*p*_^*2*^ = 0.05). The effect over rOT is almost entirely driven (97%) by the difference between the body odor and the two other odor contexts (*F* (1, 24) = 11.4, *p* = .003, = 0.32). Moreover, individual *Z*-scores over rOT revealed that the facelike-selective response is significant for 10 participants out of 26 in the baseline and gasoline odor contexts, increasing up to 17 participants in the body odor context. Over lOT, significant individual responses are observed for only 4 (baseline), 6 (body), and 7 (gasoline) participants out of 26.

For the face-selective response (Figure 3, left panel), amplitude varies between 2.86 ± 0.29 and 2.91 ± 0.32 µV across odor contexts over rOT. Over lOT, the response is of 2.35 ± 0.19 and 2.30 ± 0.25 µV in the baseline and gasoline odor contexts, respectively, while slightly lower in the body odor context (2.09 ± 0.24 µV), as indicated by a marginal *Odor* × *Hemisphere* interaction (*F* (2, 48) = 2.49, *p* = .093, *η*_*p*_^*2*^ = 0.09). The car-selective response (Figure 3, middle panel) is even more stable across odor contexts, its amplitude varying between 1.60 ± 0.18 and 1.63 ± 0.21 µV over rOT, and between 1.13 ± 0.10 and 1.19 ± 0.11 µV over lOT.

### Body odor effect on the facelike-selective response according to reported face pareidolia

Figure 4 depicts the facelike-selective response differentiated between participants according to their reported awareness of facelike objects post-stimulation. Interestingly, the body odor effect previously described over the right hemisphere for the whole group of participants appears more clearly visible for perceptually aware than unaware participants. Albeit slightly increased by the body odor, the amplitude of the facelike-selective response is close in the three odor contexts for unaware participants (body: 0.39 ± 0.11 µV, gasoline: 0.33 ± 0.14 µV, baseline: 0.34 ± 0.12 µV), while more strongly increased by body odor for aware participants (body: 0.62 ± 0.13 µV, gasoline: 0.35 ± 0.18 µV, baseline: 0.14 ± 0.11 µV; about 153% larger amplitude in the body odor context; Figure 4A).

**Figure 4.**
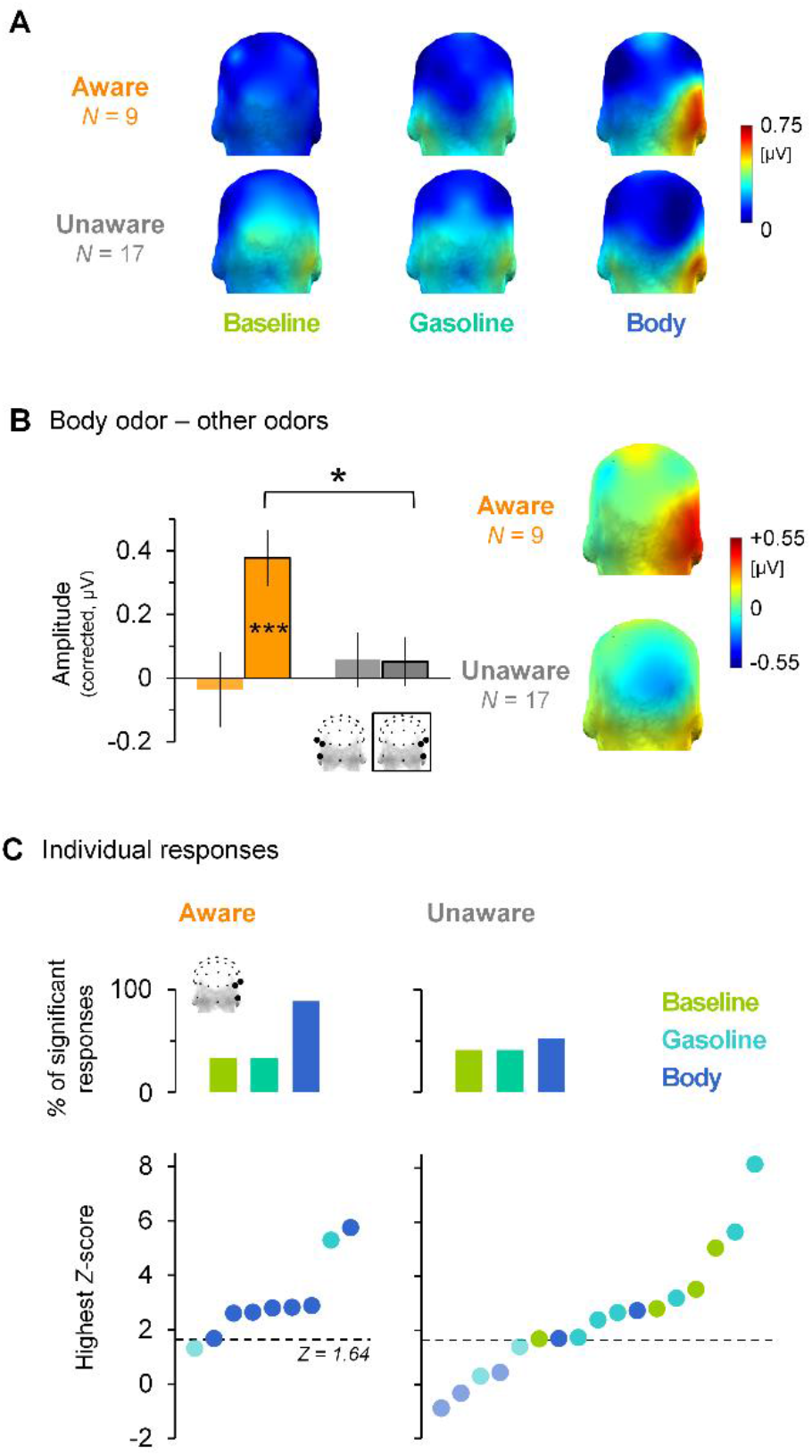
Facelike-selective response according to odor context and perceptual awareness of facelike objects. **A**. Summed baseline-corrected amplitudes of the facelike-selective response for each group of participants (perceptually aware and unaware) and odor context (green: baseline odor, light blue: gasoline odor, dark blue: body odor) illustrated by 3D head-maps (back view). **B**. Body odor effect (body minus other odors) for each group of participants (aware: orange, unaware: grey) and hemisphere (lOT: unframed, rOT: framed) with corresponding 3D head-maps (back view). Error bars represent standard errors of the mean, *: *p* < .05; ***: *p* < .001. **C**. Significance of individual responses over rOT. Top: Proportion of significant individual responses by odor context for perceptually aware (left) and unaware (right) participants. Bottom: highest individual *Z*-scores (each dot represents a participant) ranked in ascending order for each group of participants and colored by odor context (green: baseline odor, light blue: gasoline odor, dark blue: body odor). Significance threshold (*Z* > 1.64, *p* < .05) is indicated by the dotted line (non-significant *Z*-scores are lighter).

Hence, to further investigate the difference between the body odor context and the two other contexts (i.e., the body odor effect) on the facelike-selective response for both groups of participants, we calculated the amplitude difference between the body odor and the mean of the two other odors (Figure 4B) and conducted another ANOVA using *Hemisphere* (rOT, lOT) as a within-subject factor and *Group* (aware, unaware) as a between-subject factor. This analysis yielded a non-significant effect of *Group* (*F* (1,24) = 1.59, *p* = .22, *η*_*p*_^*2*^ = 0.06) and a marginal effect of *Hemisphere* (*F* (1,24) = 4.15, *p* = .053, *η*_*p*_^*2*^ = 0.15), qualified by a significant *Hemisphere* × *Group* interaction (*F* (1,24) = 4.38, *p* = .047, *η*_*p*_^*2*^ = 0.15). The body odor effect is larger for perceptually aware than unaware participants in the right hemisphere (aware: +0.38 ± 0.09 µV vs. unaware: +0.05 ± 0.08 µV; *F* (1,24) = 6.81, *p* = .015, *η*_*p*_^*2*^ = 0.22), but not in the left (aware: -0.04 ± 0.12 µV vs. unaware: +0.06 ± 0.09 µV; *F* < 1). Therefore, the body odor effect is larger in the right than the left hemisphere only for aware participants (*F* (1,24) = 6.52, *p* = .017, *η*_*p*_^*2*^ = 0.21; unaware: *F* < 1). *Z*-scores calculated on the mean body odor effect for each group of participants additionally showed that the effect is significant only for perceptually aware participants in the right hemisphere (rOT: *Z* = 3.81, *p* < .001 vs. lOT: *Z* = -0.25, *p* = .80; unaware participants: rOT and lOT: *Z* = 0.58 and 0.51 respectively, all *p*s >.56).

Finally, individual responses are markedly different across odor contexts according to participants’ awareness of facelike objects. *Z*-scores calculated for each odor context over rOT revealed that only 3 out of 9 perceptually aware participants (i.e., 33%) have a significant facelike-selective response in the baseline and gasoline odor contexts (*Z* > 1.64, *p* < .05), compared to 8 participants (89%) in the body odor context (Figure 4C). By contrast, the number of perceptually unaware participants with a significant response only slightly increases from 7 out of 17 (41%) in both baseline and gasoline contexts to 9 (53%) in the body odor context. In addition, when considering only participants with at least one significant response in one odor context (i.e., 8 and 12 participants for the aware and unaware groups, respectively), 7 (88%) aware participants have their highest *Z*-score in the body odor context compared to only 2 (17%) unaware participants (Figure 4C).

### General visual response

The 12-Hz streams of images elicit a clear neural activity at the same frequency and harmonics over the middle occipital cortex, reflecting the general visual response to all cues rapidly changing 12 times per second (e.g., local contrast). Summed across significant harmonics (Figure 5), the response has a mean baseline-corrected amplitude of 1.83 ± 0.16 µV, and is very robust, with every single participant having a significant response over the middle-occipital ROI (*Z*-scores ranging from 28 to 228 when collapsing the nine conditions, all *p*s < 0.001). The general visual response does not differ as a function of the visual category displayed at 1.33 Hz, the odor context or the perceptual awareness of facelike objects (all *F*s < 2.07, all *p*s > .13).

**Figure 5.**
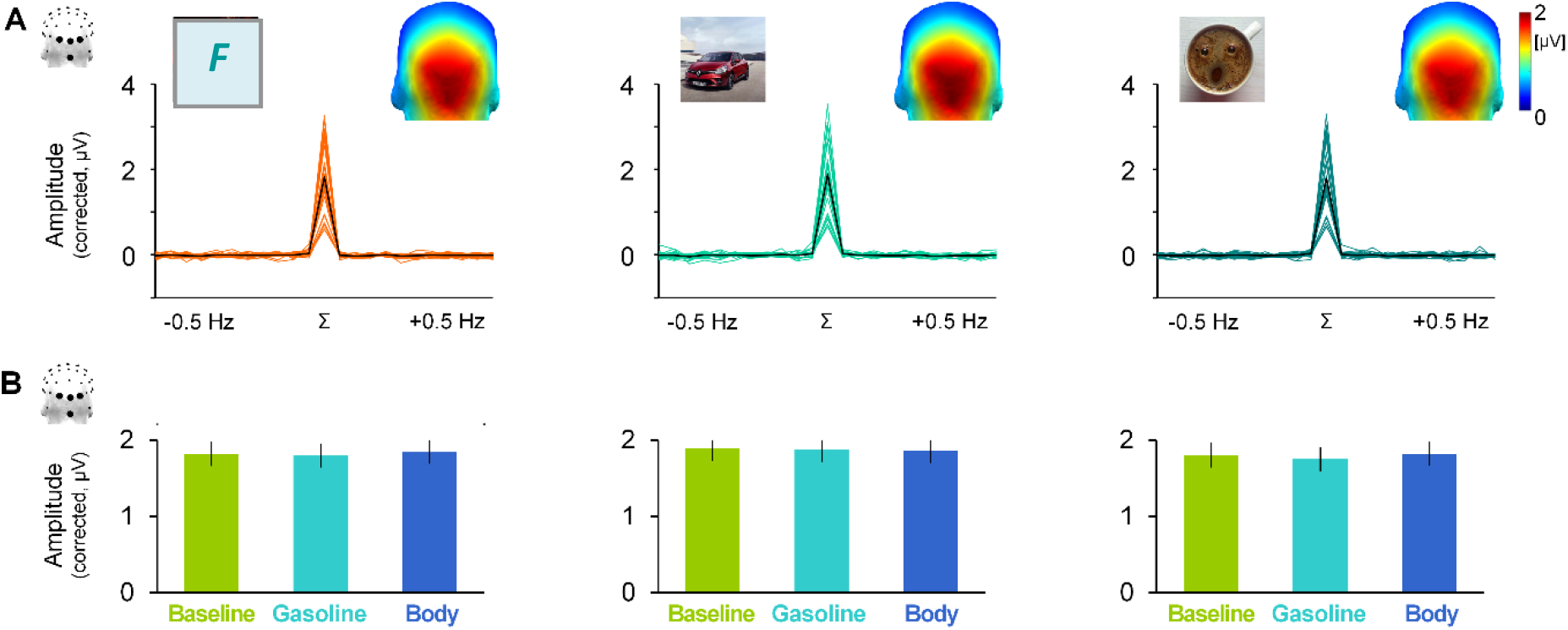
General response to the rapid visual stream. **A**. Baseline-corrected amplitude of the general visual response summed across significant harmonics (Σ) compared to surrounding frequencies (± 0.5 Hz, baseline-corrected amplitude ≈ 0, signal ≈ noise) over the middle occipital ROI (O1, Oz, O2, Iz) for each category averaged across odor contexts. Individual spectra are depicted by colored lines and the mean amplitude by a black line, together with corresponding 3D head-maps (back view). **B**. Summed baseline-corrected amplitude of the general response dissociated by odor context (green: baseline odor, light blue: gasoline odor, dark blue: body odor for each visual category. Error bars represent standard errors of the mean.

## Discussion

By using EEG frequency-tagging to track categorical occipito-temporal responses to faces, cars and facelike objects, and by implicitly exposing participants to body, gasoline, and baseline odor contexts, we provide evidence for the influence of congruent, but not incongruent, odors on rapid and automatic visual categorization at both group and individual levels. This olfactory-visual interaction is effective only when the target category is ambiguous, i.e., body odor selectively facilitates the neural categorization of a variety of facelike objects as faces, especially for participants who report their presence. No odor effect is observed on the middle occipital response elicited by the fast train of stimuli, or on the behavioral response to the cross-detection task, ruling out any general influence of the mere presence of odors. The present study thus reveals a facilitating effect of congruent odors on neural visual categorization when the interpretation of the visual input is equivocal, in line with the disambiguating function of multisensory integration (Ernst and Bülthoff, 2004).

Following prior studies using the same approach (e.g., Jacques et al., 2016a; Rossion et al., 2015), we provide a direct measure of neural visual categorization in the form of category-selective responses (i.e., differential responses to the target categories relatively to numerous and diversified other living and non-living objects) that generalize across variable category exemplars. These responses reflect rapid (i.e., each image appears for 83 ms) and automatic categorization (i.e., visual stimuli were irrelevant to the explicit cross-detection task). The two unambiguous categories (i.e., faces and cars) elicit a robust selective response at the predefined 1.33-Hz frequency and harmonics, clearly visible in the amplitude spectrum and highly reliable across individuals. The facelike-selective response is also clearly isolated in the EEG frequency spectrum, albeit less reliable across participants. The magnitude of the category-selective responses differs across categories, with the largest response for human faces and the lowest response for facelike objects, corroborating previous studies (Hagen et al., 2020; Jacques et al., 2016a; Rekow et al., 2021a). Every category-selective response is mostly distributed over the occipito-temporal cortex with a right hemisphere advantage, in line with the critical role of this region in automatic visual categorization (Bugatus et al., 2017; Gao et al., 2018; Hagen et al., 2020; Jacques et al., 2016b).

Importantly for our purpose, we found that the selective response to facelike objects over the right hemisphere is about two times larger in the presence of body odor. This observation accords well with previous studies showing that congruent odors modulate visual object perception at both behavioral (e.g., Robinson et al., 2013; Seigneuric et al., 2010; Seo et al., 2010; Zhou et al., 2010) and neural (e.g., Ohla et al., 2018) levels. In addition, it indicates that body odors, as powerful “social chemosignals” conveying much information about our conspecifics (de Groot et al., 2017 for review), do not only influence the perception of fine-grained facial information (e.g., facial expression; Adolph et al., 2013; Mujica-Parodi et al., 2009; Rubin et al., 2012; Wudarczyk et al., 2016; Zernecke et al., 2011; Zhou and Chen, 2009), but also improve the generic categorization of a visual stimulus as a face. It is worth noting that neither a facilitating nor an inhibiting odor effect was observed for incongruent associations (i.e., gasoline effect for face/facelike stimuli or body odor effect for cars), contrary to a recent report of incongruent odors interfering with visual categorization (Hörberg et al., 2020). However, in that latter study, interference was observed on explicit behavior and late (i.e., 300-900 ms) frontal and parietal EEG responses to a delayed auditory cue signaling that the visual stimulus must be categorized. Thus, odors could have not interfered with the automatic visual categorization of the stimulus at its onset, but rather with the controlled and delayed decision on that stimulus.

Our study reveals that odors are specifically prone to facilitate the visual categorization of congruent inputs when their perceptual interpretation is not straightforward, i.e., for facelike objects. Indeed, genuine human faces are unambiguously categorized from the sole visual input even under high stimulation constraints. At brief durations (i.e., 83 ms as in the present study; Retter et al., 2020), or with degraded inputs (i.e., low-pass filtered stimuli; Quek et al., 2018), all participants report having seen faces and the face-selective response measured with EEG frequency-tagging is already saturated (and remains stable when presentation conditions become less constraining). By contrast, even with full-spectrum stimuli presented for a longer duration (i.e., 167 ms), not all participants notice facelike objects, which elicit a lower category-selective response than human faces (Rekow et al., 2021a). This is likely due to the fact that facelike stimuli represent various objects similar to the other objects displayed in the stimulation sequence (see Figure 1). Therefore, the visual system must go beyond this similarity to produce a differential response to facelike vs. other objects and generalize this response across widely variable facelike objects. In this situation, inputs from other sensory systems are ideal to resolve ambiguity according to prior multisensory experience (Ernst and Bülthoff, 2004). Because body odors are often associated with faces and heighten attention to person-related cues (Cecchetto et al., 2019), they are well-suited to tilt the balance towards the “face” interpretation. This is consistent with previous studies showing that odors disambiguate facial expression perception (Forscher and Li, 2012; Leleu et al., 2015a; Mujica-Parodi et al., 2009; Novak et al., 2015; Rubin et al., 2012; Zernecke et al., 2011; Zhou and Chen, 2009), and more broadly with the inverse effectiveness principle, whereby multisensory integration is particularly effective when the response to unisensory stimuli is scarce (e.g., Stein and Meredith, 1993; Regenbogen et al., 2016).

Such inverse relationship between olfactory-visual integration and the strength of the sole visual response has already been observed for facelike categorization in infants (Rekow et al., 2021b). At the neural level, the disambiguating effect of odors suggests effective connectivity between the olfactory and the visual systems, in line with body odors activating the lateral fusiform gyrus (Prehn-Kristensen et al., 2009; Zheng et al., 2018; Zhou and Chen, 2009), a category-selective visual region that hosts face-selective areas. It was also observed that the sole presentation of odors activates the right occipital cortex (Djordjevic et al., 2005; Royet et al., 2001, 1999; Zatorre et al., 2000), suggesting that odors alone can trigger visual imagery (Parma et al., 2017 for review). In sum, odors could function as a prime to improve the detection of congruent inputs in other sensory modalities, e.g., body odors alerting to the potential presence of a person, thus in the present case, favoring the categorization of a face in common objects configured as faces.

Regarding hemispheric asymmetry, the body odor effect on facelike categorization is confined to the right hemisphere, and there is a non-significant decrease of the face-selective response with body odor over the left hemisphere. In fact, these observations relate to previously reported modulations of both face- and facelike-selective neural responses in infants exposed to maternal body odor (Leleu et al., 2020; Rekow et al., 2021b, 2020). This indicates that body odor reinforces the well-known dominance of the right hemisphere for face perception (Behrmann and Plaut, 2020; Grill-Spector et al., 2017; Hagen et al., 2020; Jonas et al., 2016). This right-hemispheric dominance has been related to the perception of the whole face configuration (Caharel et al., 2013; Rossion et al., 2011). Hence, we can speculate that body odor, by evoking the presence of a person, favors the perception of a whole face pattern from a single fixation at a stimulus interpretable as facelike. In addition, systematic reviews on the hemispheric lateralization of the neural responses to odors proposed that the right hemisphere is more involved than the left in the recognition of the odor source (Brand et al., 2001; Royet, 2004). Therefore, the right hemisphere also appears as a good candidate for integrating information across the senses to facilitate the categorization of (multi)sensory inputs, putatively relying on large-scale connectivity between distant brain regions dedicated to the same semantic domain (Mahon and Caramazza, 2011).

Strikingly, the body odor effect on facelike categorization is mainly observed for perceptually aware participants, i.e., participants who report face pareidolia post-stimulation. One may thus suggest that body odor, by enhancing facelike-selective neural activity in the visual cortex, triggers the subjective experience of a face in facelike objects. Admittedly, awareness status was defined after the experiment based on a single declarative report. It is thus unclear whether body odor made some participants become aware of the facelike objects, or whether the odor effect is observed because participants were already aware of the facelike objects. However, two elements lead us to favor the first interpretation. First, the magnitude of category-selective responses measured with EEG frequency-tagging has been previously related to participants’ awareness of the visual category (facelike objects: Rekow et al., 2021a; human faces: Retter et al., 2020), with larger amplitudes when participants explicitly report perception. By contrast, here, the facelike-selective response is of close amplitude for perceptually aware and unaware participants in the baseline and gasoline odor contexts (and even slightly lower for aware participants in the baseline context). This suggests that aware participants were not more generally prone to face pareidolia than unaware participants, but specifically more sensitive to facelikeness when exposed to body odor. Second, in a side experiment, we tested another 26 participants for their ability to report face pareidolia after being presented with similar 12-Hz visual streams of facelike vs. nonface objects without any odor context (see Supplementary Information for details). Only 4 participants (15%) noticed the presence of facelike objects compared to 9 aware participants (35%) in the main experiment. This observation thus converges with the interpretation that exposition to body odor elicits awareness of facelike objects in some participants. Future studies should obviously elaborate on this issue by directly manipulating awareness in a single group of participants.

In conclusion, we show that neural visual categorization - i.e., the ability of the brain to rapidly and automatically respond to a given visual category - is shaped by concurrent odor inputs, provided they are congruent (i.e., semantically-related) with the visual stimulus and can facilitate its interpretation (disambiguation). It is worth noting that while our results indicate a specific association between body odor and facelike categorization, we cannot exclude the same type of association for other categories, including cars. Actually, we rather consider that the influence of odors on visual categorization is a general phenomenon when the visual information is ambiguous - odors orienting perception towards the most probable visual category (Ernst and Bülthoff, 2004). Here we used a large set of facelike objects as ambiguous faces, without any equivalent for cars. Indeed, face pareidolia is ubiquitous in humans and pareidolia less often occurs for nonface visual categories, reflecting the high saliency of the “face” category for our species. Future studies should thus evaluate the generalizability of our findings to various visual categories by using degraded stimuli or challenging stimulation parameters that hamper visual categorization.

## CRediT roles

**Diane Rekow**: Conceptualization, Investigation, Data curation, Formal analysis, Methodology, Visualization, Writing - original draft, Writing - review & editing. **Jean-Yves Baudouin**: Conceptualization, Funding acquisition, Project administration, Writing - review & editing. **Karine Durand:** Funding acquisition, Writing - review & editing. **Arnaud Leleu**: Conceptualization, Methodology, Funding acquisition, Project administration, Formal analysis, Supervision, Writing - review & editing.

## Acknowledgment

This work was financially supported by the “Conseil Régional Bourgogne Franche-Comté” (PARI grant), the FEDER (European Funding for Regional Economic Development), the French “Investissements d’Avenir” program, project ISITE-BFC (contract ANR-15-IDEX-03; to JYB, KD and DR), and the French National Research Agency (contract ANR-19-CE28-0009 to AL). The authors are grateful to Renaud Brochard and the “Institut Universitaire de France” for financial support in performing EEG, to body odor donors, and to every participant of the pilot and main studies.

## Notes

### Competing Interest Statement

The authors have declared no competing interest.

