## Supplement for "Smell what you hardly see: Odors assist categorization in the human visual cortex"

bioRχiv
THE PREPRINT SERVER FOR BIOLOGY

PREPRINT

### Smell what you hardly see: Odors assist categorization in the human visual cortex

Diane Rekow, Jean-Yves Baudouin, Karine Durand, Arnaud Leleu

*Supplementary information*

#### Supplementary methods

##### Collection and selection of odor stimuli

###### Body odor collection

The collected body odor corresponds to axillary sweat. Volunteer donors complied to a strict 24-hour hygiene procedure forbidding the use of odorous products (soap, perfume) on the chest and armpits as well as the consumption of specific food and substances (marinated or spicy food, garlic and onion, tobacco and alcohol) before collection. On the day of collection, they washed their armpit with clear water and a cotton glove (pre-washed with scentless powder detergent (Persavon, France)) at the lab. A trained experimenter then disposed cotton pads under each armpit using dedicated and disposable scentless nitrile gloves and fixed the pads with adhesive strip on the original paper sack of the sterile cotton pads. Collection lasted between 45 and 150 minutes while donors participated in a non-related study. Cotton pads were then removed with gloves, and cut in 16 equivalent units, before being immediately stored in tinfoil and an individual zip-locked plastic bag in a -20°C freezer. They were used within 6 months to preserve odor characteristics (Lenochova et al., 2008). All donors and participants of the EEG experiment were of Caucasian origin.

###### Independent characterization of body odor pools

In a first pilot experiment, body odors of 8 individuals (4 females, mean age: 27 ± 5 years-old; mean sampling duration: 60 ± 10 min) were gathered as a single pool (not used in the main EEG experiment). Twelve independent participants (7 females, mean age ± SD: 30 ± 8 years old) rated twice this body odor pool *vs.* two unworn cotton pads soaked with distilled water in same-looking flasks presented under a counterbalanced order across participants. Participants rated the cotton pads on 1–9 Likert scales for their pleasantness (1: not pleasant at all, 9: very pleasant), their intensity (1: barely perceptible, 9: very intense) and familiarity (1: not familiar at all, 9: very familiar). The mean hedonic valence of the body odor pool was 3.75 ± 1.08 (vs. 4.63 ± 0.8 for water: $t_{11} = 2.22$, $p = .05$) for a perceived intensity of 4.25 ± 1.85 (vs. 2.83 ± 1.86 for water: $t_{11} = 2.20$, $p = .05$). Odor conditions were judged of similar familiarity (body: 3.29 ± 2.18, water: 3.54 ± 1.75; $t_{11} = 0.74$, $p = .34$).

**Body odor pools for the EEG experiment**

The body odor of 16 novel independent non-smoker donors (8 females, mean age ± *SD*: 25 ± 4 years old) were collected after they followed the 24-hour hygiene procedure (see above). Two pools comprising the samples of 8 individuals each (4 females) were created by matching sampling duration and age across pools (Table S1; all *t*s < 1.95, all *p*s > .07). Each participant of the EEG experiment was exposed to only one pool (N = 13 for each pool). Importantly, each group of participants (depending on their overt report of facelike objects, i.e., perceptually aware *vs.* unaware participants) was equally presented with both odor pools (aware: 5 and 4 participants for pool 1 and 2, respectively; unaware: 8 and 9 participants for pool 1 and 2, respectively).

**Table S1. Details of the body odor pools used in the EEG experiment.** From the 16 body odor samples, two pools of 8 individual odors were composed in adjusting for sex ratio (1:1), sampling duration and age of donor.

| | | Sampling duration (min) | Age of donor (year) |
|---|---|---|---|
| All | Range | 45–150 | 20–35 |
| | Mean ± SD | 92 ± 34 | 25 ± 4 |
| Pool 1 | Range | 45–150 | 22–35 |
| | Mean ± SD | 98 ± 41 | 27 ± 5 |
| Pool 2 | Range | 45–120 | 20–27 |
| | Mean ± SD | 87 ± 28 | 23 ± 3 |

**Selecting and adjusting the gasoline odor**

Based on a previous study in which 38 independent participants (22 females, mean age ± *SD*: 25 ± 4 years old) rated 50 odors from flasks (Vieillard et al., 2020), we identified that the odor of gasoline matches body odor pools in terms of hedonic valence (mean ± *SD*: 3.76 ± 1.7 on a 1–9 Likert scale, when using a $10^{-3}$ dilution in mineral oil). We thus decided to retain a gasoline odor with $10^{-2}$ and $10^{-3}$ dilutions in mineral oil for a direct comparison with the body odor pools used in the main EEG experiment. In this second pilot experiment, fourteen independent participants (10 females, mean age ± *SD*: 21 ± 2 years old) evaluated 4 odorants delivered with the diffusing system of the EEG cabin (see Materials and Methods for details). The odorants consisted in the two body odor pools and the two concentrations of gasoline. Each odorant was diffused 4 times in a randomized order (i.e., corresponding to a total of 16 trials) avoiding immediate repetition during 5 seconds each and with a minimum inter-stimulus interval of 15 seconds. Participants had to rate odorants on 1–9 Likert scales for their pleasantness (1: not pleasant at all, 9: very pleasant), their familiarity (1: not familiar at all, 9: very familiar) and their intensity (1: barely perceptible, 2: very intense) immediately after the 5 seconds of diffusion. Messages on the screen indicated whether participants had to smell or rate what they previously smelled. The $10^{-3}$ diluted gasoline evoked similar intensity (2.63 ± 1.88), pleasantness (5.09 ± 0.81) and familiarity (3.84 ± 2.06) than both body odor pools (means: 2.54 ± 1.30, 4.95 ± 0.69 and 3.87 ± 1.60, respectively; all *t*s < 0.9, all *p*s > .05), unlike the $10^{-2}$ dilution judged more intense, more pleasant and more familiar (means: 4.16 ± 2.11, 5.80 ± 1.99 and 5.95 ± 2.37, respectively; all *t*s > 2.82, all *p*s < .01). The $10^{-3}$ dilution of gasoline was thus chosen for the main experiment.

**Post-EEG odor ratings**

After the experimenter disclosed the implicit diffusion of odors, participants were blindly presented with the flasks of the 3 samples used during their own EEG session. They were asked to rate the odorants on 1–9 Likert scales in the same order they had been exposed to. Ratings included pleasantness (1: not pleasant at all, 9: very pleasant), familiarity (1: not familiar at all, 9: very familiar) and intensity (1: barely perceptible, 9: very intense). For the whole sample of participants, gasoline and body odors did not differ in perceived pleasantness ($t_{25}$ = 1.95, $p$ = .06), intensity ($t_{25}$ = 0.41, $p$ = .68) or familiarity ($t_{25}$ = 1.05, $p$ = .30). Furthermore, dissociating perceptually aware and unaware participants revealed no difference in ratings (all *t*s < 1.6, all *p*s > .12). The full post-test rating scores are available in Table S2.

Additionally, participants were invited to write down any *evocation* the odor induced, considering that *identification* is hardly achieved in humans and that we used barely detectable odors. Thus, providing an answer was strongly encouraged but not mandatory. The complete set of given answers (translated from French) is compiled in Table S3. Even if some participants found the accurate identification, responses are varied and seem unrelated to the odor effect observed at the brain level. Indeed, while participants perceptually aware of facelike objects have the strongest body odor effect, their personal odor evocations are not different from the other group: among the 4 hits for body odor identifications (i.e., correct designation as "sweat"), 2 of them correspond to aware participants (i.e., 50%). Participants were equally prone to give a tentative answer (i.e., at least 1 tentative answer for the 3 odors), with 8/9 aware vs. 13/17 unaware participants (mean number of answers: 1.77 ± 1 vs. 1.83 ± 0.8, respectively; $t_{19}$ = 0.18, $p$ = .86). In addition, no difference in the accurate identification arises from descriptive group analysis (12% vs. 15% correct answers for aware and unaware, respectively; $t_{19}$ = 0.31, $p$ = .76).

**Table S2. Post-EEG odor ratings.** Participants were presented with the flasks used during their EEG session after being informed that odors have been diffused during the EEG experiment. They were asked to smell each flask (see text for details) and rate its content on 1–9 Likert scales regarding pleasantness (1: not at all to 9: very pleasant), intensity (1: barely perceptible to 9: very strong), and familiarity (1: not familiar at all to 9: highly familiar). Mean ratings ± *SD*s are presented below.

| | | **Pleasantness** | | | **Intensity** | | | **Familiarity** | | |
|---|---|---|---|---|---|---|---|---|---|---|
| | *N* | **Baseline** | **Body** | **Gasoline** | **Baseline** | **Body** | **Gasoline** | **Baseline** | **Body** | **Gasoline** |
| **All** | ***26*** | **4.9 ± 1.8** | **3.7 ± 2** | **4.7 ± 1.7** | **3 ± 1.9** | **4.2 ± 2.6** | **4.5 ± 2** | **3.5 ± 2.5** | **4.2 ± 2.8** | **5 ± 2.3** |
| aware | *9* | 4.8 ± 0.7 | 3.4 ± 0.6 | 4.1 ± 0.4 | 2.8 ± 0.7 | 3.1 ± 0.6 | 4.1 ± 0.6 | 3.7 ± 0.8 | 4 ± 1 | 4.3 ± 0.7 |
| unaware | *17* | 4.9 ± 0.4 | 3.7 ± 0.5 | 4.9 ± 0.5 | 3 ± 0.5 | 4.8 ± 0.7 | 4.7 ± 0.5 | 3.5 ± 0.6 | 4.2 ± 0.7 | 5.3 ± 0.6 |

**Table S3. Post-EEG odor descriptions.** Along with the odor ratings, participants were invited to note down whatever the odor evoked. Answers were translated from French (all participants were native French speakers). Participants reporting awareness of facelike stimuli are identified by a star.

| | Baseline | Body | Gasoline |
|---|---|---|---|
| #01* | | | gasoline or permanent marker |
| #02 | | | gasoline |
| #03* | food (hazelnut) | sweat | paint |
| #04 | food (chocolate) | sweat | gasoline |
| #05* | food (banana) | | |
| #06 | | food (tortilla) | gasoline |
| #07* | | food (garlic) | plastic or playdough |
| #08 | | | food (citruses) |
| #09 | sweat | | citrus |
| #10 | | sweat | bleach |
| #11* | | | |
| #12 | | | |
| #13 | | | gasoline |
| #14* | | pencil | plastic |
| #15 | | | |
| #16 | | | |
| #17* | food (vanilla) or medicine | cloth | cleaning product (detergent) or paint |
| #18 | cleaning product | | |
| #19* | acid | acid | |
| #20 | musty smell | fat | wood |
| #21 | cleaning product (solvent) | plant | interior of a new car |
| #22 | | cat litter | cleaning product |
| #23 | | | |
| #24* | | sweat | cleaning product |
| #25 | cleaning product (detergent) | | soda |
| #26 | | | dried grass |

**Table S4. Regions of interest (ROIs) for the category-selective response.** After summing baseline-corrected amplitudes across significant harmonics of the 1.33-Hz category-selective frequency (i.e., up to 18.67 Hz; i.e., 14th harmonic), category-selective responses were considered averaged across odor contexts to estimate the regions of interest (ROI) for further analysis according to the channels with the largest amplitude among the 64 channels. For all categories, the six highest channels were located over the right and left occipito-temporal cortices, such that two ROIs were defined (rOT and lOT, respectively in blue and red). See Figure S1 A below for electrode location.

| Faces | | | Cars | | | Facelikes | | |
|---|---|---|---|---|---|---|---|---|
| **Channel** | **Rank** | **Amplitude** | **Channel** | **Rank** | **Amplitude** | **Channel** | **Rank** | **Amplitude** |
| P10 | 1 | 3.70 | P10 | 1 | 1.86 | P10 | 1 | 0.51 |
| P9 | 2 | 2.94 | PO8 | 2 | 1.59 | P9 | 2 | 0.34 |
| P8 | 3 | 2.48 | P9 | 3 | 1.42 | P8 | 3 | 0.30 |
| PO8 | 4 | 2.46 | P8 | 4 | 1.37 | PO8 | 4 | 0.27 |
| P7 | 5 | 1.92 | PO7 | 5 | 1.09 | PO7 | 9 | 0.19 |
| PO7 | 6 | 1.88 | P7 | 6 | 0.96 | P7 | 13 | 0.16 |

**Table S5. Regions of interest (ROIs) for the general visual response.** After summing baseline-corrected amplitudes across significant harmonics of the 12-Hz image presentation frequency (i.e., up to 48 Hz; i.e., 4th harmonic), the general visual response was considered averaged across odor contexts to estimate the region of interest for further analysis according to the channels with the largest amplitude among the 64 channels. For all categories, the exact same four highest channels were located over the middle occipital (mO) cortex, constituting a single ROI. See Figure S1 B below for electrode location.

| Faces | | | Cars | | | Facelikes | | |
|---|---|---|---|---|---|---|---|---|
| **Channel** | **Rank** | **Amplitude** | **Channel** | **Rank** | **Amplitude** | **Channel** | **Rank** | **Amplitude** |
| Iz | 1 | 1.93 | Iz | 1 | 2.00 | Iz | 1 | 1.90 |
| Oz | 2 | 1.83 | Oz | 2 | 1.90 | Oz | 2 | 1.81 |
| O1 | 3 | 1.82 | O1 | 3 | 1.87 | O1 | 3 | 1.79 |
| O2 | 4 | 1.71 | O2 | 4 | 1.75 | O2 | 4 | 1.67 |

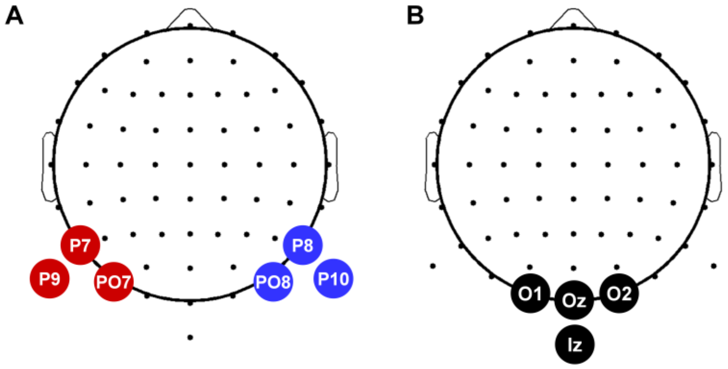


**Figure S1. 2D head-maps showing the regions of interest (ROIs) for the category-selective response (A) and the general visual response (B).** Each set of colored electrodes corresponds to one ROI. See Tables S4 and S5 for more details on (A) and (B), respectively.

#### Additional behavioral experiment

In a previous EEG frequency-tagging study (without any odor context) presenting 6-Hz stimulation sequences of nonface and facelike objects (every 6 stimuli; i.e., at 1 Hz), 48% (i.e., 13 out of 27) of the participants reported the presence of facelike objects post-stimulation (Rekow et al., 2021). In the present study, the proportion of perceptually aware participants dropped to 35% (i.e., 9 out of 26), but the stimulation rate was twice as fast (i.e., 12 Hz) with facelike objects at 1.33 Hz. Hence, we conducted a side behavioral experiment to estimate the expected proportion of aware participants when using 12-Hz sequences with facelike objects at 1.33 Hz and without any odor context. Twenty-six novel participants performed a cross-detection task while presented with 12-Hz sequences of nonface objects with facelike objects interspersed at 1.33 Hz (exact same stimuli as in the main EEG experiment). Twelve 27-second-long sequences were presented before the experimenter asked participants whether they had perceived facelike objects in the sequence. Only 15% (i.e., 4 out of 26) of the participants reported awareness of facelike objects. Hence, compared to this theoretical proportion in the absence of contextual odors, the proportion of perceptually aware and unaware participants in the main EEG experiment is significantly higher ($\chi^2_1$ = 7.39, $p$ = .007).
